# Lack of vision shifts occipital dynamics toward a frontal-like regime and enhances top-down connectivity

**DOI:** 10.64898/2026.09.18.752597

**Authors:** Gabriel Hassan, Isabella De Cuntis, Gianluca Gaglioti, Giulia Furregoni, Elena Focacci, Francesca Baglio, Davide Bottari, Giulio Bernardi, Alessandra Calcagno, Pietro Pietrini, Emiliano Ricciardi, Marcello Massimini, Mario Rosanova, Silvia Casarotto

## Abstract

The human occipital cortex is robustly repurposed for non-visual cognition after blindness, yet it remains unknown whether this functional reassignment is accompanied by a fundamental retuning of its intrinsic neural dynamics. Here, we recorded the electroencephalographic (EEG) responses to transcranial magnetic stimulation (TMS) of frontal and occipital areas to causally probe local cortical reactivity and large-scale effective connectivity in 16 blind individuals and 16 sighted controls. Direct perturbation of the occipital cortex revealed a profound shift in its intrinsic operating regime: compared with sighted participants, blind individuals exhibited faster, lower-amplitude TMS-evoked potentials that closely resembled the electrophysiological signature of the frontal cortex, which turned out to be comparable in the two populations. At the network level, source analyses further showed a specific enhancement of frontal-to-occipital effective connectivity, whereas occipital-to-frontal propagation remained unchanged. Importantly, among blind participants, faster occipital dynamics were associated with stronger frontal-driven occipital recruitment, directly linking local temporal retuning to enhanced top-down network influence. Together, these findings indicate that blindness reshapes both the intrinsic dynamics and large-scale integration of the occipital cortex. We propose that successful functional reassignment of the occipital cortex requires not only changes in cortical connectivity but also a tuning of the temporal operating regime toward a faster dynamics, enabling its embedding into distributed cognitive networks and providing a mechanistic framework for the functional reassignment of deafferented cortex.

## INTRODUCTION

Permanent lack of vision induces structural (Paré et al. 2023) and functional (Fine and Park 2018; Ricciardi et al. 2020) changes in visual pathways. Structural remodeling primarily affects the deafferented retino-geniculate-striate pathways, whereas functionally the occipital cortex is consistently engaged by non-visual perceptual and cognitive tasks. Together, these changes are compatible with experience-dependent reweighting of existing circuits and computational functions rather than a simple sensory takeover (Ricciardi et al. 2014; Makin and Krakauer 2023; Park and Fine 2024). In accordance with this functional repurposing, repetitive transcranial magnetic stimulation (TMS) of the occipital cortex can disrupt verbal processing and Braille reading in blind individuals (Amedi et al. 2004; Kupers et al. 2007). In addition, single-pulse TMS may elicit somatotopically organized tactile sensations in the fingers of expert Braille readers (Ptito et al. 2008), suggesting an involvement of the occipital cortex in non-visual information processing. At the electrophysiological level, blindness is associated with attenuation of posterior alpha rhythm (8-13 Hz) and with a shift toward higher-frequency activity relative to sighted individuals at eyes-closed (Kriegseis et al. 2006; Schepers et al. 2012; Lubinus et al. 2021; Ossandón et al. 2023). However, spontaneous EEG recordings do not fully uncover the specific neural mechanisms behind these spectral differences and in blind individuals the occipital cortex cannot be directly probed through its canonical sensory pathway. Therefore it remains unknown to what extent the functional repurposing of the occipital cortex is related to changes in its intrinsic input–output dynamics and to specific large-scale connectivity patterns.

TMS combined with EEG (TMS-EEG) represents the ideal tool to directly address this question, since it can challenge cortical circuits independently of sensory input (Ilmoniemi et al. 1997) and reveal intrinsic input-output properties not accessible from spontaneous activity alone (D’Ambrosio et al. 2023; Casarotto et al. 2024). TMS-evoked potentials (TEPs) offer a perturbational readout of the local properties of cortical circuits (Rosanova et al. 2009; Ferrarelli et al. 2012; Vallesi et al. 2021), which represents a direct way to investigate how blindness tunes the intrinsic dynamics of the occipital cortex, normally assigned to vision-related functions. At the whole brain level, TMS-EEG can probe both feedforward and feedback effective connectivity, offering a causal and directional perspective on cortical dynamics (Paus et al., 2001; Mattavelli et al., 2013; Morishima et al., 2009; Casali et al., 2010) and providing a means to test whether blindness shapes the directional embedding of the occipital cortex within large-scale networks.

In this work, we applied TMS-EEG to directly probe the reactivity of the occipital cortex in blind individuals, relative to sighted controls. By targeting the superior occipital gyrus bilaterally and the left superior frontal gyrus, we assessed both local plastic changes in intrinsic cortical reactivity and rearrangements in long-range effective connectivity along the anterior–posterior axis. We observed a substantial retuning of the temporal regime of the occipital cortex in blind individuals, which shifted toward typical frontal dynamics. In addition, we found a specific strengthening of frontal-to-occipital effective connectivity that correlated with the acceleration of the occipital regime, suggesting that visual deprivation shapes both the local dynamics and large-scale integration of the occipital cortex within anterior associative networks.

## RESULTS

We enrolled sixteen blind individuals (5 females; age = 52.8 ± 9.8 years, mean ± SD; Table S1) with complete lack of vision due to pre-chiasmatic impairments, either since birth (n = 9) or later in life (n = 7). First, we characterized the common electrophysiological features of blindness by comparing blind individuals altogether with a group of 16 sighted participants (8 females; age = 51.1 ± 8.6 years; Table S1). Then, we investigated the potential effect of blindness onset by comparing between early and late blind individuals. We recorded spontaneous EEG to establish whether the spectral characteristics of background activity in blind individuals were more comparable to sighted individuals at eyes open (EO) or at eyes closed (EC). We then collected TMS-evoked potentials by stimulating the superior occipital gyrus bilaterally and the left superior frontal gyrus, to probe posterior sites normally involved in visual information processing as well as anterior sites responsible for executive and cognitive functions. Morphological and spectral features of TEPs were compared between groups and further contextualized through the estimation of effective connectivity between frontal and occipital regions at the source level.

### 1. Spontaneous EEG

Topographic maps of resting-state power spectral density (PSD) were compared between groups and conditions (Fig. 1A). In line with the previous literature, sighted individuals showed significantly higher PSD in the α-band (8-13 Hz) at EC than at EO, without differences in the other frequency ranges (Fig. 1B, *EC-EO*). When comparing blind and sighted individuals, significant differences emerged only in the EC condition, characterized by higher α-band power across most electrodes (Fig. 1B, *EC-Blind*). Notably, this difference was no longer present when considering sighted individuals at EO (Fig. 1B, *EO-Blind*). No other significant differences emerged between blind and sighted individuals in the low β (14-20 Hz) and high β (21-30 Hz) ranges.

**Fig. 1.**
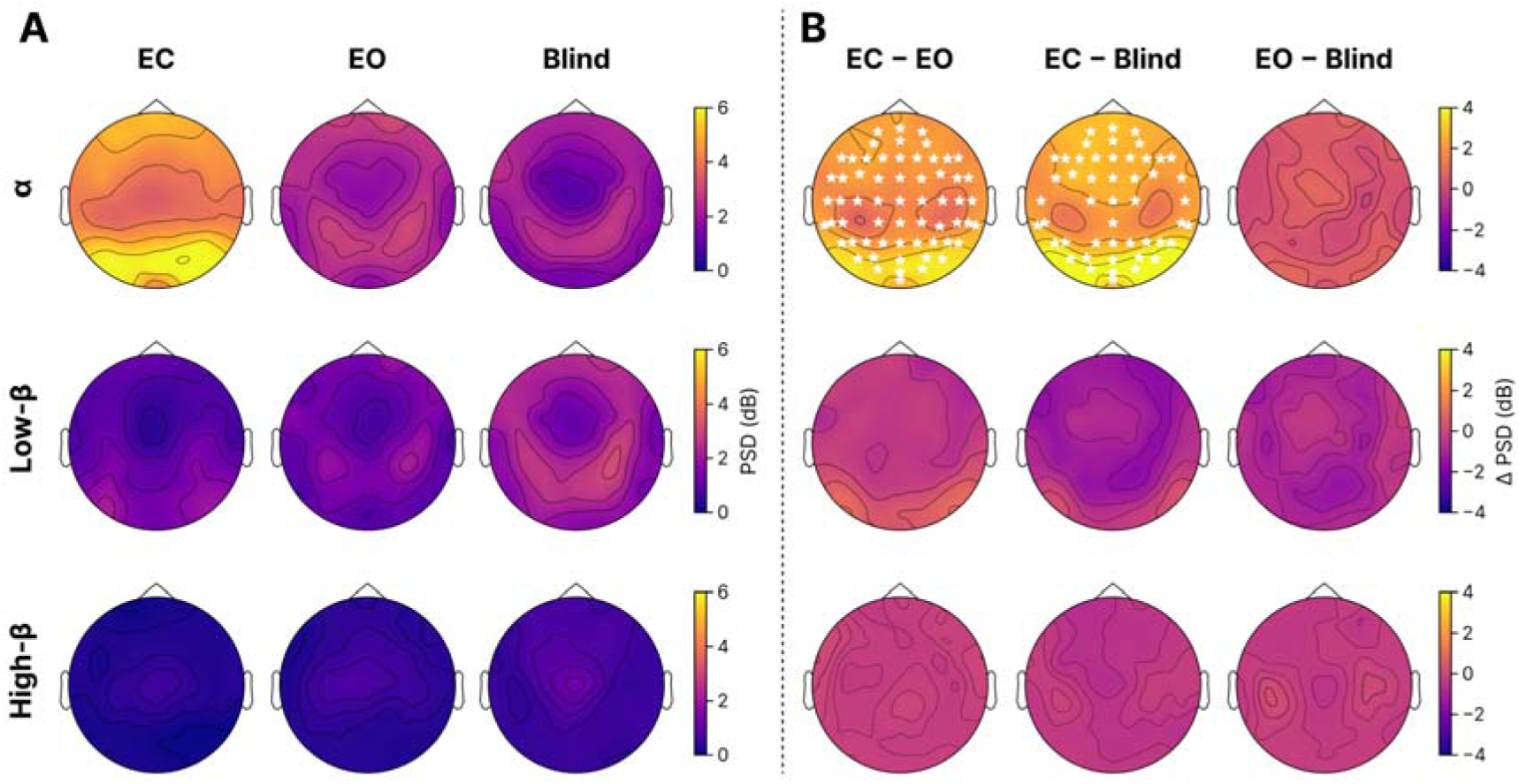
Spectral characteristics of background EEG activity. **(A)** Topographical distribution of grand-average power spectral density (PSD) values in the α (8-13 Hz), low-β (14-20 Hz), and high-β (21-30 Hz) bands for sighted individuals at eyes-closed (*EC*), sighted individuals at eyes-open (*EO*), and blind individuals (*Blind*). **(B)** Grand-average pairwise PSD differences between sighted individuals at EC and at EO (*EC-EO*, Wilcoxon signed rank test), between sighted individuals at EC and blind individuals (*EC-Blind*, Mann–Whitney U test) and between sighted individuals at EO and blind individuals (*EO-Blind*, Mann–Whitney U test). White stars indicate the electrodes with corrected-p < 0.05 (see Data analysis: spontaneous EEG paragraph in the Materials and Methods section). Significant differences were observed only in the α band for the comparison between *EC* and both *EO* and *Blind*. No significant differences were observed between *EO* and *Blind*.

### 2. TMS-evoked potentials

We evaluated TEP features of blind individuals in comparison to sighted individuals at EO, since the spectral characteristics of spontaneous EEG in blind individuals were more comparable to sighted individuals at EO than at EC. We ascertained that stimulation parameters across the three cortical regions were comparable between blind and sighted individuals in terms of TMS target coordinates (see TMS parameters paragraph in the Material and Methods section) and estimated intensity of the induced electric field (Fig. S1).

We focused sensor-level analysis of TEPs on morphological and spectral features computed on 4 region-of-interest (ROI) channels close to the stimulation site, following a semi-automatic procedure described in Hassan, Gaglioti et al. (2026). Briefly, we extracted: *(i)* **A_P1-P2_**, the peak-to-peak amplitude of the first waveform component, indexing the gain of local cortical recruitment following the TMS pulse (Fig. 2A); *(ii)* **IPI**, the inter-peak interval of the first oscillatory cycle, derived from the time lag between successive TEP components (Fig. 2B); *(iii)* **NF**, the natural frequency defined as the frequency bin with largest post-stimulus evoked power (Fig. 2C). Further details about the computation of these features are reported in the Materials and methods section (Data analysis: TEPs at the sensors level paragraph).

**Figure 2.**
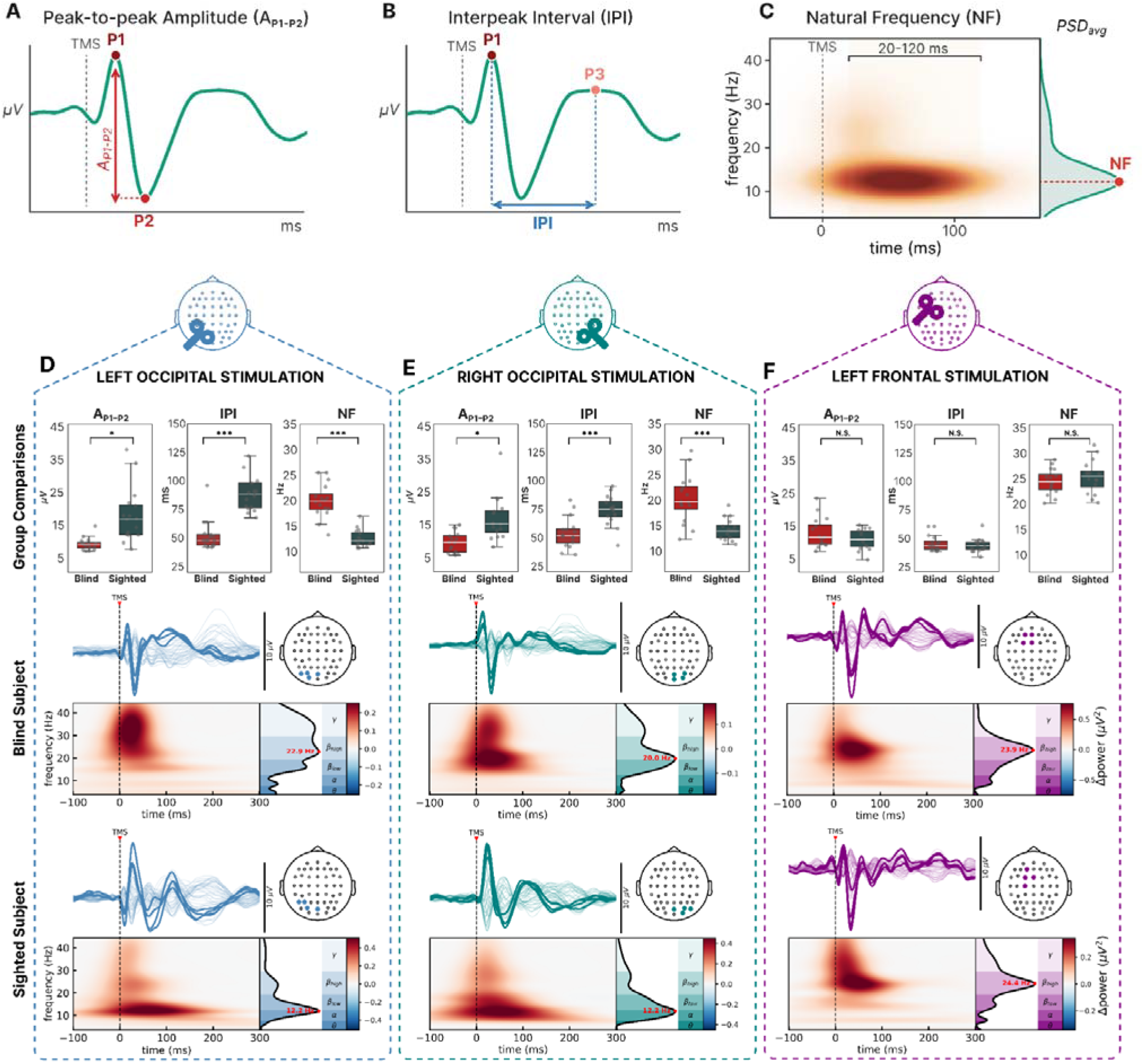
Morphological and spectral TEP features across cortical targets in blind and sighted individuals. (A–C) Schematic illustration of the three principal local TEP measures. A_P1–P2_ (A) is the absolute amplitude difference between P1 and the subsequent opposite-polarity P2. The interpeak interval (IPI; B) is the latency between P1 and the subsequent same-polarity P3. Natural frequency (NF; C) is the frequency corresponding to the maximum post-stimulus power spectral density cumulated between 20 and 120 ms (PSD_avg_). (D–F) Results following left occipital (D), right occipital (E), and left frontal (F) stimulation. Group-level boxplots compare blind and sighted participants for A_P1–P2_, IPI, and NF. Bilateral occipital stimulation resulted in lower A_P1–P2_, shorter IPI, and higher NF in blind than in sighted participants (D,E), whereas no significant group differences were observed following frontal stimulation (F; * corrected P < 0.05, *** corrected P < 0.001, N.S. corrected P > 0.05; Mann–Whitney U tests). Butterfly plots show all-channel average TEPs from representative blind (middle row) and sighted eyes-open participants (bottom row). Bold traces correspond to the filled colored dots in the scalp topographies and identify the channels near the stimulation site automatically selected for computing morphological and spectral features. Below each butterfly plot, baseline-corrected time–frequency power is shown together with PSD_avg_, obtained by accumulating post-stimulus power between 20 and 120 ms; NF is indicated in red as the frequency bin with maximum evoked power.

#### 2.1 Occipital TEPs exhibit reduced amplitude and faster dynamics in blind compared to sighted individuals

The comparison between blind and sighted individuals for each stimulation site separately showed that morphological and spectral TEP features significantly differed only following occipital stimulation. Specifically, A_P1-P2_ was markedly lower in blind than sighted individuals (median values: left occipital 8.43 μV *vs.* 16.20 μV; right occipital 8.88 μV *vs.* 14.49 μV; Fig. 2D,E top row, Tab S2), indicating a reduced local cortical recruitment following stimulation. Blind participants also exhibited a shorter inter-peak interval (median values: left occipital 48 ms *vs* 88.5 ms; right occipital 51.5 ms *vs.* 75.0 ms; Fig. 2D-E top row, Tab. S2), reflecting a faster temporal succession of TEP components. Finally, natural frequency was about 1.5 times higher in blind than sighted participants (median values: left occipital 20.0 Hz *vs.* 12.2 Hz; right occipital 19.7 Hz *vs.* 14 Hz; Fig. 2D,E top row, Tab. S2), indicating a shift toward faster intrinsic oscillatory dynamics. In contrast, frontal TEPs showed comparable morphological and spectral features across groups (Fig. 2F top row, Tab. S2). The same pattern of occipital TEP differences was observed when blind were compared to sighted individuals in the EC condition, thus indicating that these effects are independent of the transient sensory and oscillatory state associated with visual input, and instead reflect stable neurophysiological adaptations to blindness (Fig. S2). Representative TEPs from one blind and one sighted participant are shown in Fig. 2 D,F (Blind-middle row; Sighted - bottom row). Within the blind group, the comparison of morphological and spectral TEP features between early-onset and late-onset individuals did not result in any significant difference (Fig. S3).

#### 2.2 Posterior-to-anterior difference of cortical dynamics is reduced in blind individuals

When frontal and occipital TEPs were compared within each group, sighted participants exhibited the expected posterior-to-anterior acceleration of cortical dynamics (Rosanova et al. 2009). Frontal TEPs showed higher NF, shorter IPI, and lower A_P1–P2_ amplitude than occipital TEPs bilaterally (Fig. S4A,C,E; Table S2). The fronto-occipital amplitude pattern was reversed in blind individuals, with slightly but significantly smaller A_P1–P2_ values following stimulation of either occipital cortex than frontal cortex (Fig. S4B). Notably, the induced electric field was higher at the occipital sites, suggesting that the reduced amplitude of the occipital response was not attributable to weaker stimulation (Fig. S1). Moreover, in blind individuals IPI did not differ between frontal and bilateral occipital sites (Fig. S4D), whereas frontal NF was higher than left, but not right, occipital NF (Fig. S4F). Thus, occipital dynamics in blind individuals approached those observed over the frontal cortex, although the extent of this convergence differed between hemispheres.

To further investigate the thinning of fronto-occipital differences in blind individuals, the left and right occipital TEPs of each participant were assigned, according to NF, to a relatively “slower” and a relatively “faster” cluster of evoked responses. Both TEP clusters in blind participants showed higher NF (Fig. 3A) and shorter IPI (Fig. 3D) than their counterparts in sighted controls. Even the slower TEP cluster in blind participants exhibited significantly higher NF (Fig. 3A) and shorter IPI (Fig. 3D) values than the faster TEP cluster in sighted controls, indicating that the shift toward faster dynamics was not driven by a subset of participants. Moreover, in blind individuals the faster TEP cluster did not significantly differ from left frontal TEPs, neither in NF nor in IPI values (Fig. 3B,E), and their grand-average TEP waveforms showed a similar overall temporal profile (Fig. 3G). By contrast, in sighted participants the typical fronto-occipital differentiation remained clearly preserved (Fig. 3C,F), as also qualitatively evident from the superimposition of the respective grand-average TEPs (Fig. 3H)

**Figure 3.**
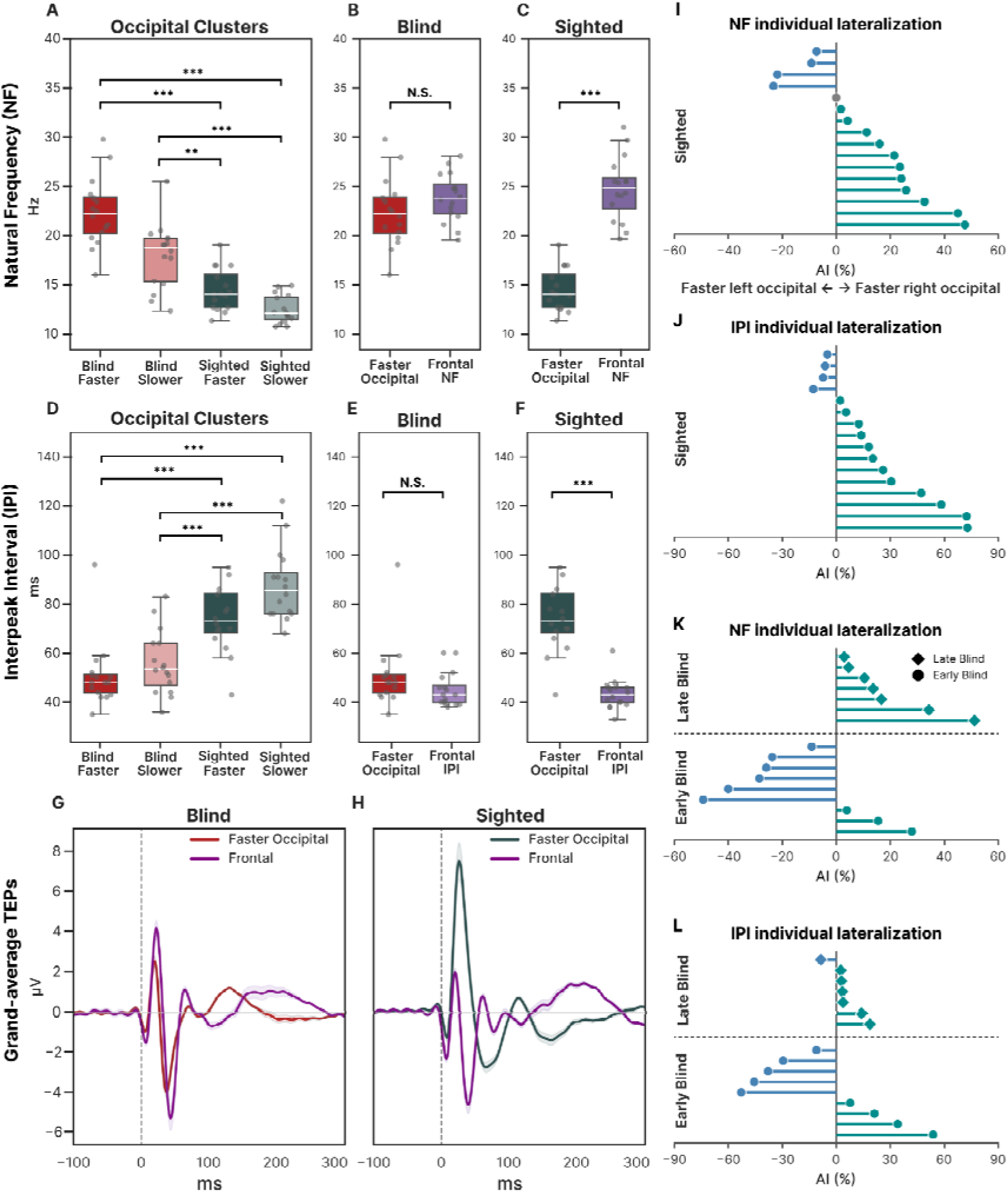
Occipital TEP time scales, cross-area comparisons and hemispheric lateralization. (A) Occipital responses stratified into faster and slower clusters according to NF. Both clusters showed higher NF in blind than sighted participants, and the slower blind cluster exceeded the faster sighted cluster. (B,C) Paired comparisons between faster occipital and frontal NF in blind (B) and sighted (C) participants, showing loss of occipital–frontal frequency differentiation in blindness. (D–F) Corresponding analyses of IPI, showing a complementary pattern. (G,H) Grand-average TEP waveforms for the faster occipital cluster and frontal stimulation in blind (G) and sighted (H) participants. (I–L) Individual signed occipital asymmetry indices, calculated as (AI(%)=(X_R-X_L)/(X_R+X_L) x 200), for NF (I,K) and IPI (J,L) in sighted participants (I,J) and in early- and late-onset blind participants (K,L). Late-onset blind participants showed a consistent rightward NF lateralization. Boxplots indicate the median and interquartile range; dots and lollipops represent individual participants. Between-group comparisons used Mann–Whitney U tests and paired comparisons used Wilcoxon signed-rank tests, with FDR-BY correction. (*p<0.05), (**p<0.01), (***p<0.001); N.S., not significant.

#### 2.3 The interhemispheric organization of occipital dynamics changes with blindness onset

To determine whether the occipital responses assigned to the faster and slower clusters were preferentially associated with one hemisphere, we compared TEP features between left and right occipital stimulation. In sighted participants, A_P1–P2_ was comparable between hemispheres (Fig. S4A), whereas IPI was significantly shorter in right occipital TEPs (Fig. S4C). NF was also slightly higher on the right, although this difference did not reach statistical significance (Fig. S4E). In blind participants, no significant interhemispheric differences were found for A_P1–P2_, IPI or NF (Fig. S4B,D,F).

Since group-level comparisons may conceal individual asymmetries occurring in opposite directions, we then calculated an Asymmetry Index (AI; Isaias et al. 2016) for NF and IPI as *AI(%)=(X_R-X_L)/(X_R+X_L)×200*, where *X* represents either NF or IPI. Individual values showed that NF was higher in the right hemisphere in 11 of 16 sighted participants, with equal values in one participant, whereas IPI was higher on the right in 12 of 16 participants (Fig. 3I,J). A different pattern emerged when blind participants were separated according to blindness onset. All late-onset blind participants showed higher NF values in right than left occipital TEPs (Fig. 3K), indicating that the faster occipital cluster consistently corresponded to the right hemisphere, similar to sighted individuals. IPI was also higher on the right in 6 of 7 late-onset participants (Fig. 3L). Conversely, early blind participants showed heterogeneous asymmetries in both measures. Importantly, these onset-related differences were specific to the relative organization of the two hemispheres, as early- and late-blind participants did not differ in spontaneous spectral power (Fig. S5) or in TEP features examined separately within each hemisphere (Fig. S2).

#### 2.4 Frontal-to-occipital effective connectivity is enhanced in blind individuals

We performed source-level analysis to compare effective connectivity between groups. We estimated cortical activity by exploiting realistic head models based on individual T1-weighted MRIs and by applying the sLORETA inverse solution (Pascual-Marqui 2002) to scalp potentials. Then, we masked cortical activations that did not survive statistical comparison between pre- and post–stimulus activity (see Data analysis: Evoked activity at the source level paragraph in the Materials and methods section) and we obtained a binarized spatiotemporal matrix of significant activations. Last, we computed the time course of the percentage of significantly activated sources (SS_%_(t)) in three discrete cortical regions, representing the left superior frontal gyrus and the superior occipital gyrus bilaterally according to the Destrieux atlas.

SS_%_(t) averaged between 0 and 250 ms did not significantly differ between blind and sighted participants when measured (i) locally at the stimulation site (Fig. 4A,E,I, boxplots on the right), (ii) in the left superior frontal gyrus following stimulation of either occipital cortex (Fig. 4C,F, boxplots on the right), or (iii) in the superior occipital gyrus following stimulation of the contralateral occipital cortex (Fig. 4B,D, boxplots on the right). These findings indicate that local direct cortical activation, occipital-to-frontal propagation, and interhemispheric propagation between occipital cortices were comparable between groups.

**Figure 4.**
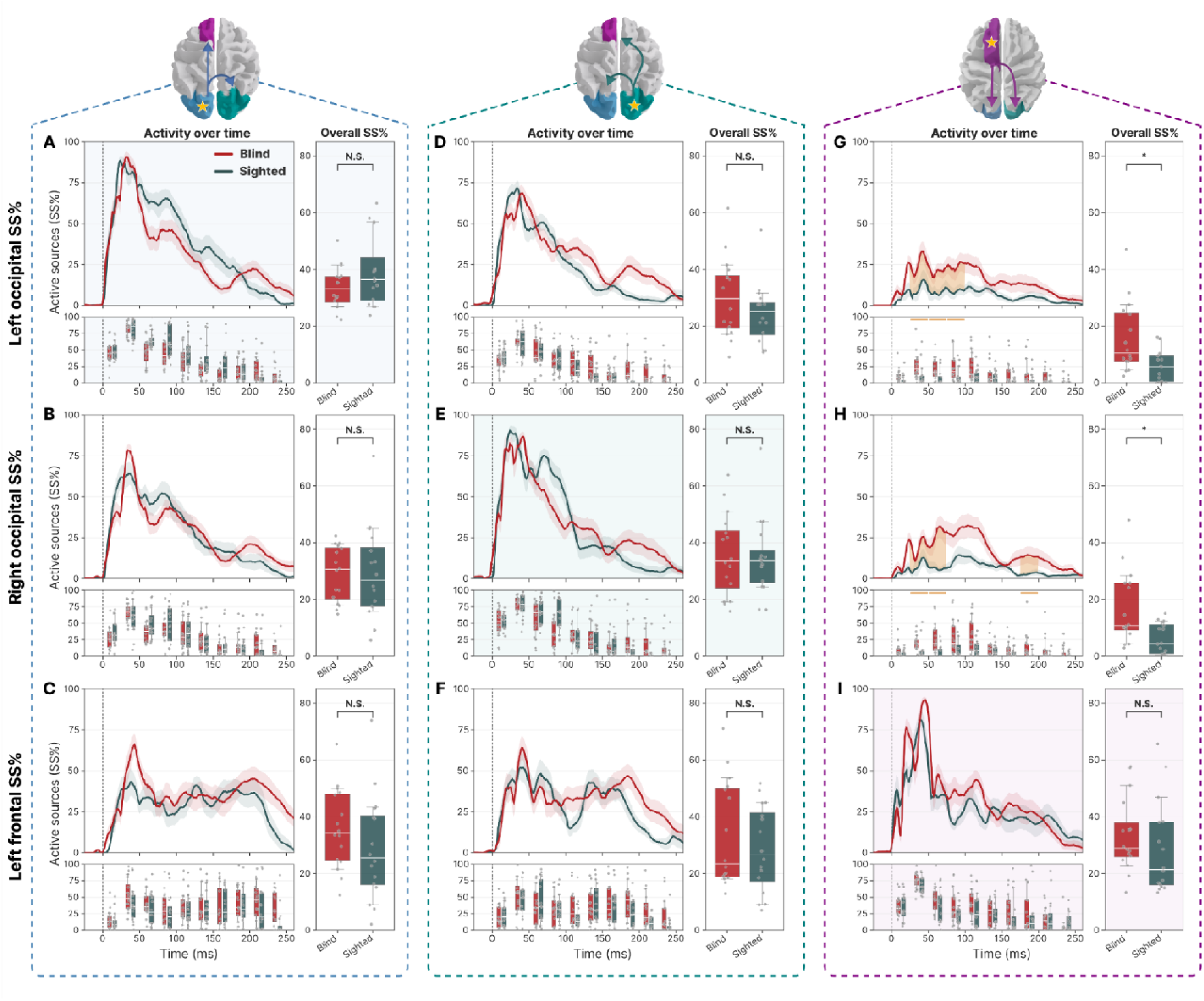
Input–output organization of TMS-evoked source activity across frontal and bilateral occipital regions. Columns show responses following stimulation of the left occipital (A–C), right occipital (D–F), and left frontal cortex (G–I), while rows represent activity measured in the left occipital, right occipital, and left frontal regions, respectively. Brain schematics indicate the stimulated site (yellow star) and propagation pathways; tinted diagonal panels identify local responses within the stimulated region. For each panel, the upper-left plot shows the time course of SS_%_ from 0 to 250 ms after TMS in blind (red) and sighted participants (dark teal; mean ± SEM). The lower-left plot shows the distributions of SS% values averaged within consecutive 25-ms bins. The right boxplot summarizes average SS_%_ across 0–250 ms. Boxes represent the median and interquartile range, whiskers the 10th–90th percentiles, and grey dots individual participants. Orange shading and markers identify bins with significant group differences based on two-sided Mann–Whitney U tests, with FDR correction across 30 comparisons per stimulation site. Significant differences were confined to bilateral occipital responses following frontal stimulation (G,H), indicating greater frontal-to-occipital recruitment in blind participants.

By contrast, frontal stimulation resulted in significantly higher occipital activation on both sides in blind compared with sighted individuals (Fig. 4G, H, boxplots on the right). Time-resolved analysis, performed by comparing SS% values averaged within consecutive 25-ms bins, localized this effect to the 25–100 ms window in the left occipital cortex and to the 25–75 ms and 175–200 ms windows in the right occipital cortex (FDR-corrected p<0.05; Fig. 4G,H). This finding indicates enhanced frontal-to-occipital effective connectivity in blind individuals, emerging as early as 25 ms post-stimulus.

#### 2.5 Faster occipital dynamics are associated with enhanced frontal-to-occipital propagation in blindness

To assess whether enhanced frontal-to-occipital propagation was associated with the retuning of occipital dynamics, we correlated mean occipital NF with bilateral occipital SS_%_ following frontal stimulation (Fig. 5). Correlations were restricted to 0–125 ms to match the post-stimulus interval used to estimate NF (20–120 ms), while retaining complete 25-ms bins. Across this window, higher occipital NF was associated with greater occipital recruitment in blind (Spearman’s ρ=0.58, *p*=0.018), but not sighted participants (ρ=0.08, *p*=0.778; Fig. 5A). Time-resolved analysis localized this association to two adjacent intervals, 50–75 ms (ρ=0.80, *corrected-p*=0.002) and 75–100 ms (ρ=0.67, *corrected-p*=0.025), in the blind group; no significant association emerged in sighted participants (Fig. 5B). Notably, the strongest association occurred within the period of maximal occipital recruitment following frontal stimulation (Fig. 4G,H). Accordingly, SS_%_ averaged across the resulting 50–100 ms interval strongly correlated with occipital NF in blind (ρ=0.77, *p*=0.0005), but not sighted individuals (ρ=0.16, *p*=0.545), with a significant difference between groups (Δρ=0.61, permutation *p*=0.0165; Fig. 5C). Moreover, the distribution of occipital P_3_ latencies largely overlapped this interval in blind participants and is shown only as a temporal reference. Together, these findings identify a temporally specific association between faster local occipital dynamics and greater frontal-driven recruitment during the period of maximal effective-connectivity reorganization in blindness.

**Figure 5.**
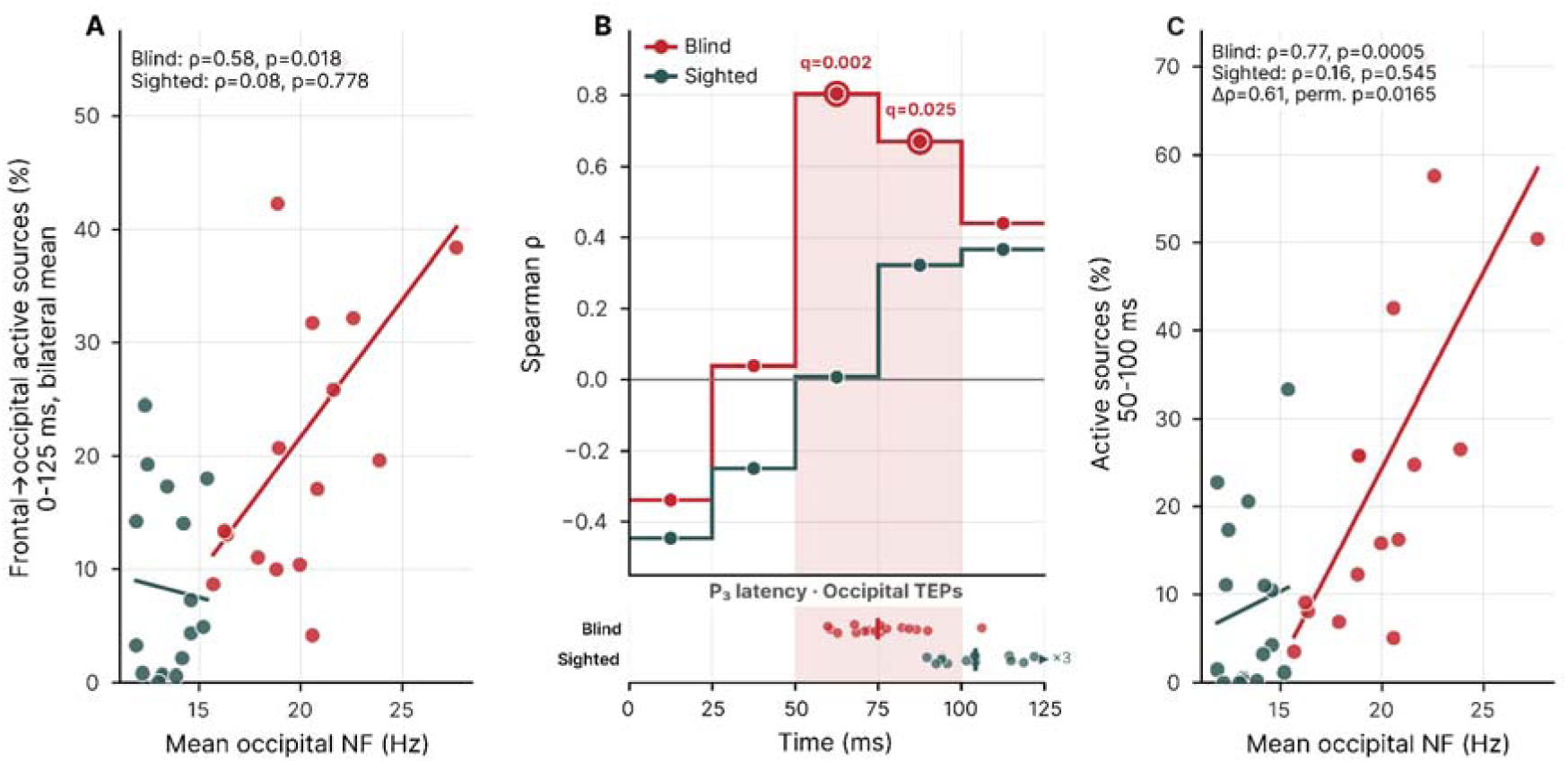
Occipital natural frequency covaries with frontal-to-occipital recruitment in blind participants. **(A)** Relationship between mean bilateral occipital NF and bilateral occipital source activity following frontal stimulation, averaged over 0–125 ms. This window encompasses the 20–120-ms interval used to estimate NF while retaining complete 25-ms bins. **(B)** Spearman correlations between occipital NF and frontal-evoked occipital activity calculated within five consecutive 25-ms bins. Open circles and colored shading identify correlations surviving FDR correction across the five bins within each group (*q*<0.05). The strip below shows the distribution of P_3_ latencies in locally evoked occipital TEPs as a temporal reference. Here, P_3_ denotes the third TEP peak; right-edge triangle indicates latencies beyond the displayed window. **(C)** Relationship between occipital NF and source activity averaged across the contiguous significant interval identified in blind participants (50–100 ms). The difference between group-specific correlations was assessed by permutation testing.

## DISCUSSION

Our study demonstrates that permanent and complete lack of retinal input - such as in congenital or late peripheral blindness - tunes the intrinsic temporal dynamics and spectral features of the occipital cortex. We show that occipital TMS-evoked potentials (TEPs) in both early and late blind individuals are faster and smaller in amplitude compared with those of sighted participants, resembling the intrinsic dynamics typically associated with frontal associative cortices. Beyond this local tuning, blindness is also associated with increased top-down effective connectivity selectively from frontal to occipital regions, revealing that lack of vision shapes both the intrinsic properties of the occipital cortex and its integration within large-scale cortical networks.

We leveraged the ability of TMS-EEG to reveal intrinsic circuit properties that may not be apparent in spontaneous activity alone (Massimini et al. 2009; Casarotto et al. 2024). This approach is particularly valuable in blindness because the occipital cortex can no longer be directly probed through its canonical sensory pathway. In our sample, resting EEG exhibited comparable occipital electrophysiological profiles between blind and sighted participants at eyes-open, indicating that the observed differences in evoked dynamics do not simply reflect altered baseline oscillatory states. Instead, these differences emerged when the occipital cortex was directly challenged: whereas stimulation of the occipital cortex in sighted participants elicited the expected response in the alpha-band, it induced faster, predominantly high-beta oscillations in blind individuals. The same pattern emerged when blind participants were compared with sighted individuals recorded at eyes-closed, further indicating that TEPs reflect stable differences in intrinsic network tuning rather than transient fluctuations of ongoing activity.

Blindness therefore appears to shift the occipital cortex toward the faster intrinsic dynamics normally associated with more anterior unimodal and transmodal regions (Capilla et al. 2022). Natural frequency (NF) and interpeak interval (IPI) converge on this interpretation, providing complementary readouts of the recurrent interactions that shape TMS-evoked responses (Cona et al., 2011; Rosanova et al., 2009; Hassan, Gaglioti et al., 2026). The marked upward shift in occipital NF suggests that the visually deprived occipital cortex operates within faster oscillatory ranges, altering the previously observed posterior-to-anterior acceleration (Rosanova et al. 2009; Vallesi et al. 2021; Capilla et al. 2022). IPI reinforces this conclusion in the time domain: together with the attenuation of early TEP components, shorter inter-peak intervals demonstrate that blindness reshapes not only the spectral composition but also the temporal architecture of occipital evoked responses.

Importantly, blindness consistently attenuated the canonical occipito-frontal acceleration pattern in brain dynamics. Across the range of occipital NF values, blind participants remained shifted toward higher frequencies relative to sighted controls, indicating that this attenuation was not confined to a small subset of strongly retuned hemispheres. At the higher end of the occipital NF distribution, the frontal–occipital gradient was no longer detectable, with occipital responses converging toward the frequency range and evoked-response profile typically observed over the frontal cortex. These findings suggest that blindness compresses the normal posterior-to-anterior organization of cortical dynamics, although the extent of this shift varies across individuals and hemispheres.

This local tuning was accompanied by a selective reorganization of effective connectivity. Neuroimaging studies have consistently shown stronger coupling between occipital cortex and anterior associative networks in blindness (Deen et al., 2015; Kanjlia et al., 2021; Anurova et al. 2015; Hasson et al. 2016; Ortiz-Terán et al. 2016). Yet whether this reorganization reflects a directional strengthening of information flow has remained unresolved. Our assessment of the effective connectivity performed at the source level addresses this question causally and directly. Frontal stimulation produced stronger propagation toward both left and right occipital cortices in blind than in sighted participants, whereas occipital-to-frontal propagation remained comparable between groups. This asymmetric pattern indicates a selective strengthening of top-down influences on visually deprived occipital cortex rather than a generalized increase in reciprocal coupling. Notably, the stimulated frontal region lies within dorsal premotor cortex, a key node of the dorsal visuomotor stream whose computations—including spatial localization, motion processing, and reach planning (Batista et al. 2007; Busan et al. 2009; Galletti and Fattori 2018)—are known to recruit occipital cortex after vision loss (Bola et al., 2023; Collignon et al., 2011; Lewis et al., 2010).

Our findings provide causal support for a long-standing hypothesis regarding the role of feedback projections in occipital plasticity. Rather than simply conveying non-visual sensory information, feedback signals from frontal and - likely - parietal associative cortices have been proposed to serve at least three complementary functions: they may provide top-down attentional and predictive signals that regulate early visual processing, recruit occipital cortex during higher-order cognitive operations, and modulate the processing of non-visual information arriving through preserved multisensory pathways (Morishima et al. 2009; Roelfsema and De Lange 2016; Fine and Park 2018; Ricciardi et al. 2020). The selective strengthening of frontal-to-occipital effective connectivity observed here provides physiological evidence that the functional weight of these influences increases after blindness. Crucially, local retuning and network reorganization were not two unrelated group-level effects: among blind participants, higher occipital NF was associated with greater bilateral occipital recruitment following frontal stimulation, with the strongest correlations between 50 and 100 ms. Thus, the individuals showing the fastest occipital dynamics were also those in whom frontal perturbation most strongly recruited the occipital cortex, directly linking local temporal tuning to top-down network influence. The same interval also overlapped with the later phase of the locally evoked occipital response, whose timing has been linked to recurrent circuit dynamics (Hassan, Gaglioti et al., 2026), consistent with broader evidence that later TMS-evoked activity incorporates recurrent corticothalamic interactions (Russo et al. 2025). Together with evidence that recurrent influences begin to shape visual cortical responses within a comparable temporal range (Lamme 1995; Lamme et al. 1999), this temporal convergence is consistent with the possibility that strengthened anterior influences interact with the recurrent dynamics of deprived occipital circuits.

Strengthened frontal-to-occipital influences may therefore progressively shape local circuits toward faster temporal scales, while the emergence of these faster dynamics may, in turn, facilitate stable communication with anterior associative networks. Although the present data cannot establish the temporal or causal order of this interaction, they support a framework in which local cortical dynamics and large-scale connectivity co-evolve in shaping the intrinsic properties of the occipital territories following visual deprivation (Plomp et al. 2014; Murray et al. 2014; Golesorkhi et al. 2021; Song et al. 2024). This perspective also provides a mechanistic framework for interpreting the heterogeneous recruitment of V1 reported across non-visual tasks (Singh et al. 2018; Ricciardi et al. 2020; Park and Fine 2024) and is in line with accumulating evidence showing that the functional organization of the blind brain often resembles that of the sighted brain during active visual processing rather than in the absence of visual stimulation (Handjaras et al. 2017; Setti et al. 2023; Orsenigo et al. 2025).

Together with the relative preservation of occipital organization, these findings suggest that visual deprivation preserves broad features of the functional scaffold while reshaping how the occipital cortex responds to inputs and interacts with distributed networks.

This process may reflect adaptation to the temporal characteristics of the inputs that increasingly engage the occipital cortex after lack of vision. Rather than becoming responsive to an entirely new class of signals, large portions of the occipital cortex already participate in multisensory processing through convergent visual–haptic, audio-visual and higher-order associative networks (Beauchamp 2005; Tal and Amedi 2009; Lacey and Sathian 2014; Murray et al. 2016; Lee Masson et al. 2016). In parallel, different sensory systems are preferentially engaged over partially distinct temporal ranges, while multisensory integration depends critically on the temporal alignment of converging inputs (Tobimatsu et al. 1999; Ross et al. 2003; Nozaradan et al. 2012; Norcia et al. 2015), underscoring the close relationship between intrinsic circuit dynamics and the temporal structure of the information being processed. Importantly, these temporal preferences appear to be shaped by experience: native sign-language experience shifts visual steady-state responses toward higher preferred driving frequencies in both deaf and hearing individuals, indicating that the temporal processing characteristics of visual cortex are not rigidly fixed by sensory modality (Stroh et al. 2022). The increasing engagement of pre-existing non-visual pathways, together with the loss of visual drive, may therefore bias occipital circuits toward dynamics better suited to the computations they increasingly support. More generally, neural oscillations adapt to the temporal and statistical regularities of sensory streams, aligning cortical excitability and prediction with the structure of incoming information (Arnal and Giraud 2012; Henry et al. 2014; Chang et al. 2018). This adaptation is unlikely to depend solely on sensory modality itself, but rather on the interaction between intrinsic developmental constraints and the temporal and statistical structure of the residual inputs that engage these circuits.

At the cellular level, this functional adaptation may be implemented through experience-dependent remodeling of local cortical microcircuits. Experimental models of sensory deprivation implicate remodeling of inhibitory circuitry, excitation–inhibition balance, and synaptic organization, all of which can alter local temporal dynamics (Gupta et al. 2000; Desgent and Ptito 2012; Richter and Gjorgjieva 2022). These changes may be accompanied by homeostatic adjustments in excitation– inhibition balance and synaptic remodeling, including microglia-mediated pruning of inhibitory synapses, which could enhance occipital responsiveness to non-visual inputs (Castaldi et al. 2020; Hashimoto et al. 2023). Although our data do not directly address these cellular mechanisms, they provide plausible substrates for the macroscopic retuning observed after blindness (Lee and Whitt 2015; Ewall et al. 2021; Makin and Krakauer 2023).

The overall temporal retuning was accompanied by onset-dependent differences in the hemispheric organization of occipital dynamics. Sighted and late-blind participants showed a modest rightward asymmetry in natural frequency, whereas no systematic lateralization emerged in early blindness. Although subgroup inferences require caution, this pattern is in accord with evidence showing that the timing of sensory deprivation may influence how occipital plasticity is spatially expressed (Fine and Park 2018; Chebat et al. 2020).

Previous studies indicate that occipital reorganization can occur following both early- and late-onset blindness, although earlier visual loss is generally associated with more robust and functionally differentiated cross-modal changes, including differences in cortical recruitment and connectivity (Voss 2019). Here, early- and late-onset blindness showed largely comparable local temporal tuning, while differing in the interhemispheric organization of occipital dynamics. This pattern is consistent with previous evidence suggesting that blindness onset may influence the degree and spatial expression of occipital plasticity (Burton et al. 2002; Burton 2003). The more pronounced onset-related differences reported in task-based or resting-state studies may therefore partly reflect differences in distributed cortical recruitment, whereas direct probing of local physiology revealed a largely shared temporal tuning across onset groups.

The current sample size and heterogeneity in blindness onset limit the strength of subgroup inferences. At the same time, this heterogeneity reflects the natural variability of the blind population and motivates larger, longitudinal studies examining how local cortical dynamics and effective connectivity emerge and interact over time (Striem-Amit 2024). Future work combining TMS-EEG with rhythmic, steady-state, or multimodal stimulation could test whether occipital tuning can be experimentally shifted and whether this is accompanied by corresponding changes in large-scale propagation.

In conclusion, blindness reshapes the occipital cortex at both local and systems levels while preserving fundamental principles of cortical organization. The deafferented visual cortex adopts faster, frontal-like intrinsic dynamics and becomes more strongly embedded within top-down associative networks, yet these changes occur within a preserved computational scaffold rather than through wholesale cortical reassignment. The association between faster occipital dynamics and greater frontal-driven recruitment further links these local and network-level changes, suggesting that temporal retuning and effective-connectivity reorganization are coupled aspects of adaptation to visual deprivation. More broadly, they suggest that cross-modal plasticity emerges from the interaction between stable developmental constraints and experience-dependent changes in network embedding, allowing preserved occipital computations to be expressed through alternative sensory channels and feedback pathways rather than through retinal input alone.

## MATERIALS AND METHODS

### Participants

We enrolled 16 individuals with permanent and complete loss of vision (5 females, age=52.8±9.8 mean±STD; Tab S1) either since birth (n = 9) or later in life (n = 7) and 16 sighted individuals as control group (8 female; age=51.1±8.6 mean±STD; Tab S1). All individuals were screened for contraindications to TMS and did not declare any relevant medical, neurological or psychiatric disorders as well as use of substances potentially affecting brain function. All the procedures conformed to the Declaration of Helsinki and were approved by the local ethics committee (Comitato Etico Milano Area 1; TMS-EEG Prot. n. 609/07/27/05/AP). All individuals signed an informed consent.

### Instrumentation

EEG was recorded with a 64-channels TMS-compatible amplifier (BrainAmp DC, Brain Products GmbH, Germany) equipped with a 10-10 montage layout cap. Bipolar electroculogram (EOG) was recorded in a diagonal montage. Additional reference and ground electrodes were located over frontal sinuses on the forehead. Input impedance was kept below 5 kΩ for all channels. Data were collected at 5000 Hz sampling rate, at 0.5 μV amplitude resolution and with a hardware filtering bandwidth from DC to 1000 Hz.

TMS pulses were delivered with a biphasic 8-coil (mean/outer winding diameter ≈ 50/70 mm, pulse duration ≈ 280 μs, focal area of the stimulation hotspot ≈ 0.68 cm²) connected to a TMS unit (Nexstim Ltd., Finland). An integrated neuronavigation software (Navigated Brain Stimulation System - NBS, Nexstim Ltd.) was used to identify stimulation targets on individual T1-weighted magnetic resonance images (MRI) collected in each subject and to estimate the intensity (in V/m) of the maximum value of the induced electric field (EF-max) on the cortex.

### Experimental protocol

Participants were seated comfortably with a headrest during the entire experimental procedure.

Spontaneous EEG (about 10 min) was recorded in all individuals at rest; in sighted individuals, data were collected during both eyes-open (EO) and eyes-closed (EC) conditions. In the EO condition, participants were instructed to fixate on a dark screen positioned approximately 1 meter away. To minimize discomfort associated with blindfolding or prolonged maintenance of a prescribed eyes-open or eyes-closed state, blind participants were allowed to rest in their preferred condition.

TMS-evoked potentials (TEPs) were collected while individuals wore in-ear earphones playing a customized noise (Russo et al. 2022) to mask the coil’s click sound. In each TMS-EEG session, we delivered between 200 and 250 stimuli with a randomly jittered inter-stimulus interval ranging from 2 to 2.3 seconds.

In all individuals, TMS-evoked potentials were recorded by targeting the left superior frontal gyrus (Brodmann’s area BA6) and the superior occipital gyrus (BA18/19) bilaterally, in random order across individuals. All sighted individuals were recorded at EO. In a subgroup of 10 sighted individuals, the occipital cortex was also stimulated at EC with the same TMS parameters, to evaluate the immediate effects of visual input deprivation on TEPs’ waveform. For each cortical site, the TMS-induced electric field (EF) was initially oriented orthogonal to the targeted gyrus with an intensity of about 100 V/m as estimated on the gyral crown by the neuronavigation software. Then, the precise location, orientation and intensity of the induced EF were adjusted to minimize scalp muscle activations and to ensure an effective stimulation of the cortex, as indicated by an early (10-50 ms) peak-to-peak amplitude of average TEPs in average reference larger than 6 μV in the channel closest to the stimulation site. This procedure was implemented by relying on a free-release software that provides a customized EEG readout in real time (Casarotto et al. 2022).

### Data Preprocessing

Spontaneous EEG data were filtered (detrend, band-pass 0.5–250 Hz, notch at 50 Hz), downsampled to 1000 Hz, and re-referenced to the average reference. Artifact-contaminated channels and discrete data segments were rejected by visual inspection. Independent component analysis (ICA) was applied to remove ocular, muscular, and ballistocardiographic artifacts. Last, rejected channels were interpolated using spherical splines.

TEPs were preprocessed as described in Casarotto et al. (2024), including pulse artifact removal (−2 to +5 ms), high-pass filtering at 1 Hz, epoching (−600 to +600 ms), manual rejection of artifact-contaminated epochs and channels, average re-referencing, ICA-based artifact removal, low-pass filtering at 45 Hz, and bad channel interpolation. See Supplementary for more details.

### Data analysis: spontaneous EEG

Power spectral density (PSD) of spontaneous EEG was estimated using Welch’s method (2 s Hamming windows, no overlap; 1-45 Hz; Gramfort, 2013). For each channel, power values were cumulated band-wise within the α (8-13 Hz), low β (14-20 Hz) and high β (21-30 Hz) ranges and then normalized by the aperiodic 1/f background obtained by fitting a linear function to the log–log power spectrum separately in a low frequency (1-20 Hz) and a high frequency (20-40 Hz) range (Colombo et al. 2019), as in Ossandón et al., 2023.

Normalized band-wise PSD values obtained in sighted individuals were compared between EO and EC conditions at each electrode using Wilcoxon signed rank tests. Between group contrasts (i.e., blind vs. sighted) were evaluated with Mann–Whitney U tests. For each pairwise comparison, correction for multiple comparisons across channels and frequency bands was performed by setting the false discovery rate bound to q = 0.05, using the Benjamini–Yekutieli procedure (Benjamini and Yekutieli 2001).

### Data analysis: TEPs at the sensors level

TEP features were computed following a semi-automatic procedure described in Hassan, Gaglioti et al. (2026). Briefly, for each EEG channel remaining after excluding the outermost lateral ones, we identified an evoked peak (P_1_) within a pre-defined time window between +10 and +40 ms after the pulse. Then, we searched for two subsequent peaks: P_2_ as the first peak following P_1_ and having an opposite polarity, and P_3_ as the first peak following P_2_ and having the same polarity as P_1_. The absolute peak-to-peak amplitude between P_1_ and P_2_ (labelled A_P1-P2_) was used to rank channels in descending order and to select a region-of-interest (ROI) of n=4 neighboring channels with the highest A_P1–P2_. In addition, we computed the interpeak interval (IPI) as the time lag between P_1_ and P_3_ in ms.

Spectral features were estimated by applying the Stockwell transform (Gaussian window width = 0.7) to single-channel average TEPs. Time–frequency power spectra between 4 and 45 Hz were then extracted for each of the 4 ROI channels. For each stimulation site, evoked power spectral density (PSD_avg_) was obtained by subtracting pre-stimulus mean power (from -500 to -100 ms) to each frequency bin and by cumulating time-frequency power values over time from +20 to +120 ms. Natural frequency (NF) was estimated as the frequency bin corresponding to the maximum of PSD_avg_.

Finally, A_P1–P2_, IPI, and NF values were averaged across the 4 ROI channels to derive synthetic features for each stimulation site and subject.

Group differences in TEP metrics (i.e., A_P1-P2_, IPI, NF) between blind and sighted individuals were assessed using pairwise Mann-Whitney U tests, corrected for multiple comparisons across all metrics using the Benjamini-Yekutieli FDR procedure (corrected-p < 0.05). Comparisons across stimulation sites for each group separately were performed with pairwise Wilcoxon signed-rank tests, corrected for multiple comparisons across all contrasts and metrics using the Benjamini-Yekutieli FDR procedure (corrected-p < 0.05).

### Data analysis: Evoked activity at the source level

We performed source-level analysis to investigate cortical effective connectivity across stimulation sites and between groups. We applied a cortical reconstruction process to individual T1-weighted MRIs using the FreeSurfer software (Fischl 2012). For each participant, we generated a three-layer boundary element model (BEM) using the watershed algorithm, with a source space comprising 5124 vertices per hemisphere (Ségonne et al. 2004). This model, integrated with digitized electrode positions, was used to build a forward solution. For each stimulation site, we estimated cortical current density values from TEPs using standardized low-resolution electromagnetic tomography (sLORETA; Pascual-Marqui 2002) with a default regularization parameter (λ² = (1/3)²). A free-orientation source model was adopted, and the norm of the dipole moment was computed at each cortical location, yielding non-negative current density values. We built the inverse operator assuming an ad hoc diagonal noise covariance matrix and accounting for the actual data rank. We applied a non-parametric bootstrap-based statistical analysis to identify the spatiotemporal dynamics of cortical currents significantly activated by TMS, following the procedure described in (Casali et al., STM2013). Briefly, each source’s activity was centralized and normalized on the mean and standard deviation of its baseline level (from -500 to -1 ms). Then, for each source, we computed surrogate baseline activity by averaging across trials after randomly shuffling pre-stimulus samples at the single-trial level and we calculated the maximum absolute value of surrogates across all sources to correct for multiple comparisons (Pantazis et al. 2005). This procedure was repeated 500 times to obtain a distribution of 500 bootstraps. A significance threshold, estimated as the one-tail 99^th^ percentile of this distribution, was applied to obtain a binary matrix SS representing the spatiotemporal distribution of significant activations: SS(x,t) = 1 if the activity of source x at time sample t is significant, = 0 otherwise. Thus, we computed a binary matrix SS with 5124 rows, corresponding to the number of sources, and 600 columns, corresponding to the post-stimulus time samples, for each subject and stimulation site. Individual MRIs were also normalized to a standard brain template for group analysis and for mapping the 3D spatial coordinates of the maximum of the induced EF (EF-max) provided by the neuronavigation software in the Montreal Neurological Institute (MNI) reference space (Chau and McIntosh 2005).

To quantify the spatial distribution of TMS-evoked cortical activity, we focused on three anatomically-defined regions corresponding to the stimulated sites: left superior frontal gyrus and superior occipital gyrus on both hemispheres segmented within the FreeSurfer software and based on the Destrieux atlas (Destrieux et al. 2010). Then, we computed the time course of the percentage of significantly activated sources within each cortical region for each stimulation target. We compared the mean significant activity over the 0–250 ms post-stimulus window, as well as over consecutive 25 ms temporal bins across the same interval between sighted and blind individuals. For each stimulation target, statistical testing used two-sided Mann-Whitney U tests with FDR correction (Benjamini– Yekutieli) to control for multiple comparisons across cortical regions and temporal bins.

## Acknowledgements

The authors are grateful to Letizia Bernardelli, Elisabetta Litterio, Marta Porro and Stefano Garzonio for their help during data acquisition. This study received funding from the University of Milan (Bando Linea 2).

## Author contributions

Conceptualization: S.C., G.H., E.R., P.P., M.R. and M.M.; Methodology: S.C., G.H., M.R.; Formal Analysis: G.H., G.G., S.C., A.C.; Investigation: S.C., G.H., G.F., E.F., I.D.C., F.B.; Data Curation: S.C., G.H., G.F., E.F., I.D.C., F.B.; Writing – Original Draft: G.H., S.C.; Writing – Review & Editing: S.C., G.H., E.R., M.M., M.R., D.B., P.P., G.B., I.D.C., A.C.; Visualization: G.H.; Supervision: S.C., G.B., E.R., M.M.; Project administration: S.C., E.R., M.M.; Funding Acquisition: S.C., M.M., P.P., E.R.

