## Supplementary materials for "Lack of vision shifts occipital dynamics toward a frontal-like regime and enhances top-down connectivity"

**Supplementary Table 1**: blind and sighted populations demographics. LE = left eye; RE = right eye.

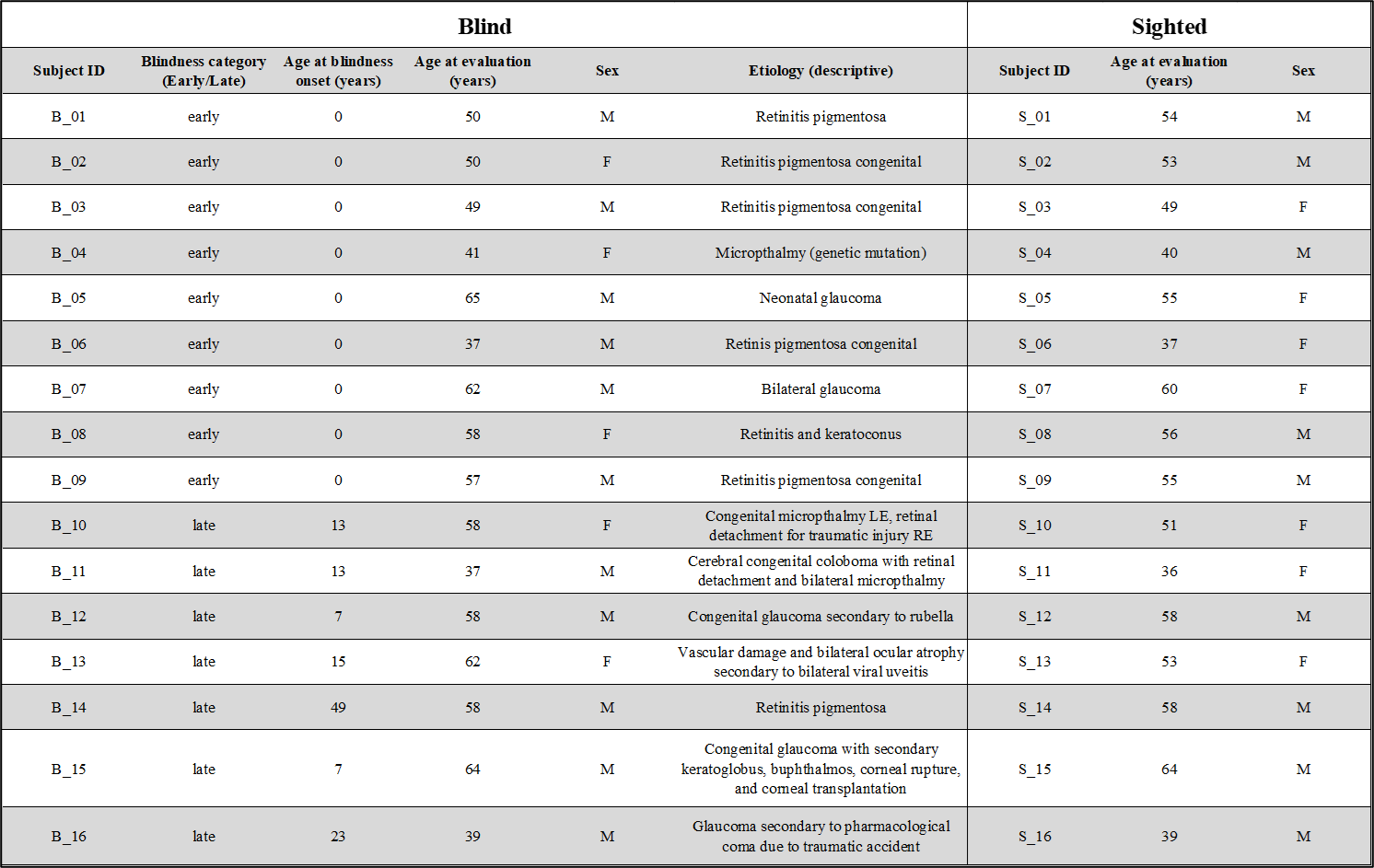

**Supplementary Table 2**: median values and IQR ranges (25-75 percentile) for each TEPs measure and group

| **Group** | **N. subjects** | **Natural Frequency (Hz)** | **A_P1-P2_ (μV)** | **Interpeak interval (ms)** |
| --- | --- | --- | --- | --- |
| Blind L Occipital | 16 | 20 (18.3-21.3) | 8.4 (7.4-9) | 48 (44-53.3) |
| Blind L Frontal | 16 | 23.9 (22.3-25.2) | 10.6 (8.6-14.4) | 43 (39.8-47) |
| Blind R Occipital | 16 | 19.8 (18.3-22.7) | 8.9 (5.9-10.9) | 51.5 (45.3-57.5) |
| Early Blind L Occipital | 9 | 20.9 (18.6-24.2) | 8.4 (7.6-8.7) | 48 (44-64) |
| Early Blind L Frontal | 9 | 23.9(22.3-24.8) | 10.6 (8.9-13.1) | 43 (40-46) |
| Early Blind R Occipital | 9 | 19 (15.1-19.7) | 7.4 (5.3-9.9) | 57 (50-70) |
| Late Blind L Occipital | 7 | 19.2 (18.1-20) | 9.5 (7.3-10.3) | 51 (46-52.5) |
| Late Blind L Frontal | 7 | 23.3 (21.6-26.9) | 10.5 (8.6-16) | 43 (40-51) |
| Late Blind R Occipital | 7 | 22 (20.5-23.1) | 9.9 (7.5-11.1) | 50 (42.5-51.5) |
| Sighted EO L Occipital | 16 | 12.2 (11.5-13.8) | 16.2 (11.5-20.5) | 88.5 (76.8-98.5) |
| Sighted EO L Frontal | 16 | 24.8 (22.7-25.9) | 9.9 (7.8-12.5) | 43 (39.8-46) |
| Sighted EO R Occipital | 16 | 14 (12.7-15.1) | 14.5 (12-18.5) | 75 (68.3-81.8) |
| Sighted EC L Occipital | 10 | 12.6 (11-13.6) | 17.3 (13.4-20.6) | 91.5 (83-101.5) |
| Sighted EC R Occipital | 10 | 12.8 (11.7-15.5) | 18.6 (14.6-23) | 81.5 (70.5-98.8) |

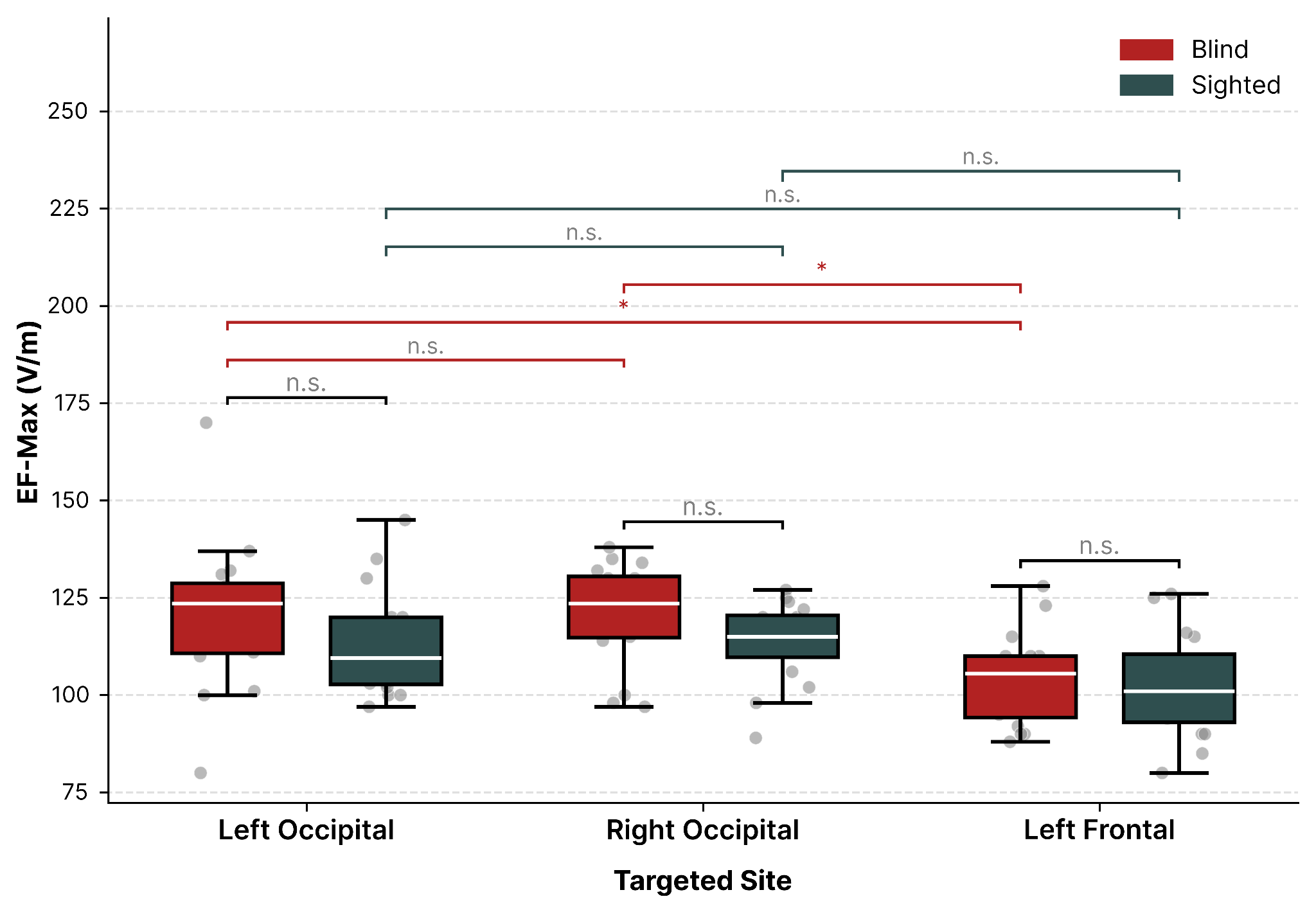

**Fig. S1. Stimulation intensity estimated by the neuronavigation software as the maximum value (in V/m) of the induced electric field (EF-max) on the cortex.**

Gray dots represent single-subject EF-max values at each targeted site. Group-level distributions are shown as boxplots (*red* for blind and *dark teal* for sighted individuals) with the boxes bounding the interquartile range divided by the median. EF-max values were not significantly different (Mann-Whitney U test, FDR corrected) between blind and sighted individuals in any targeted site. Blind participants showed significantly higher EF-max values for both occipital targets than for the frontal target, with no significant difference between the left and right occipital targets (Wilcoxon signed-rank test, FDR corrected).

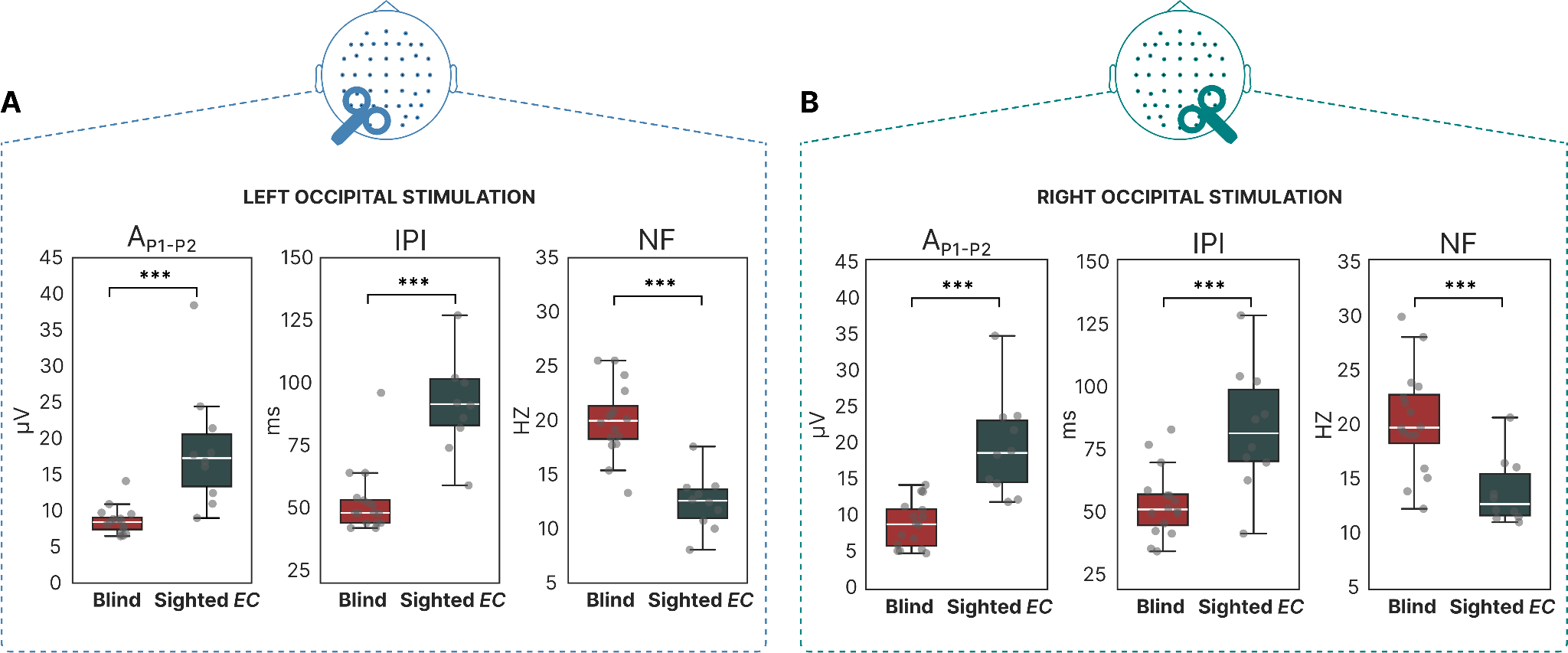

**Fig. S2.** **Comparison of morphological and spectral TEP features between blind individuals (*Blind*) and sighted individuals at EC (*Sighted EC*) following occipital stimulation.** Group-level distribution of A_P1-P2_, IPI and NF values computed on left occipital (**A**) and right occipital (**B**) TEPs. Gray dots represent single-subject values. Statistical comparisons between groups (*Blind*, red, *EC* dark teal) were performed with Mann-Whitney U tests and corrected across all contrasts and metrics using the Benjamini-Yekutieli FDR procedure (** corrected-p < .01, *** corrected-p < .001).

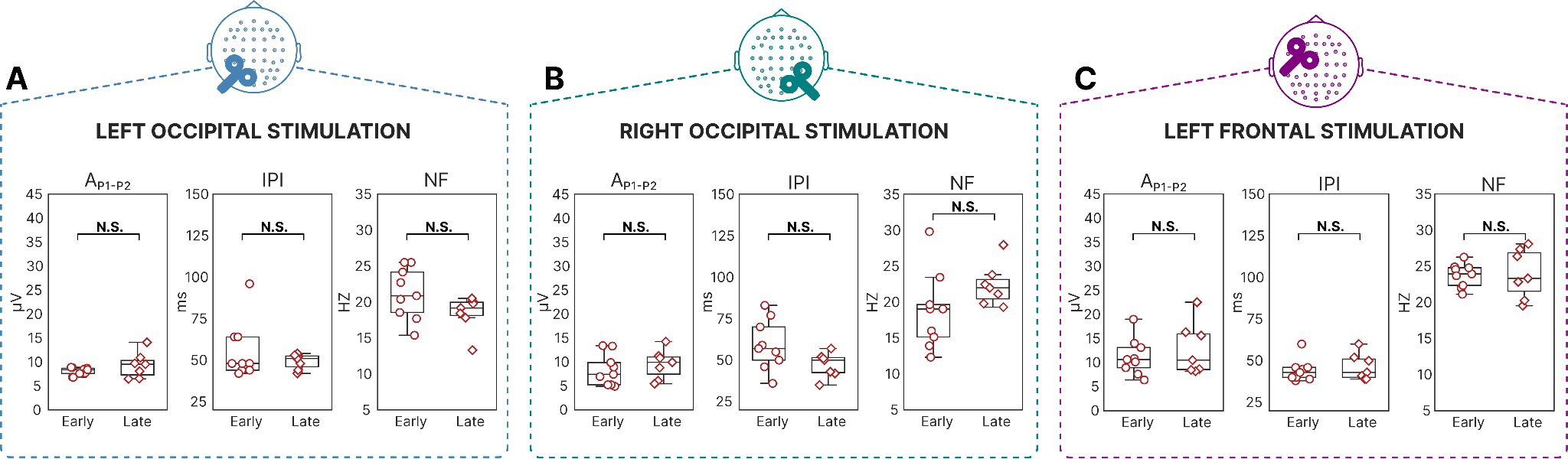

**Figure S3.** ***TEP features in early- versus late-blind participants across stimulation sites.***

Group-level comparisons of peak-to-peak amplitude (A_P1–P2_), inter-peak interval (IPI), and natural frequency (NF) between early-blind (circles) and late-blind (diamonds) participants, for Left Occipital (A), Right Occipital (B), and Left Frontal (C) stimulation. No statistically significant differences between groups were observed across any stimulation site or measure.

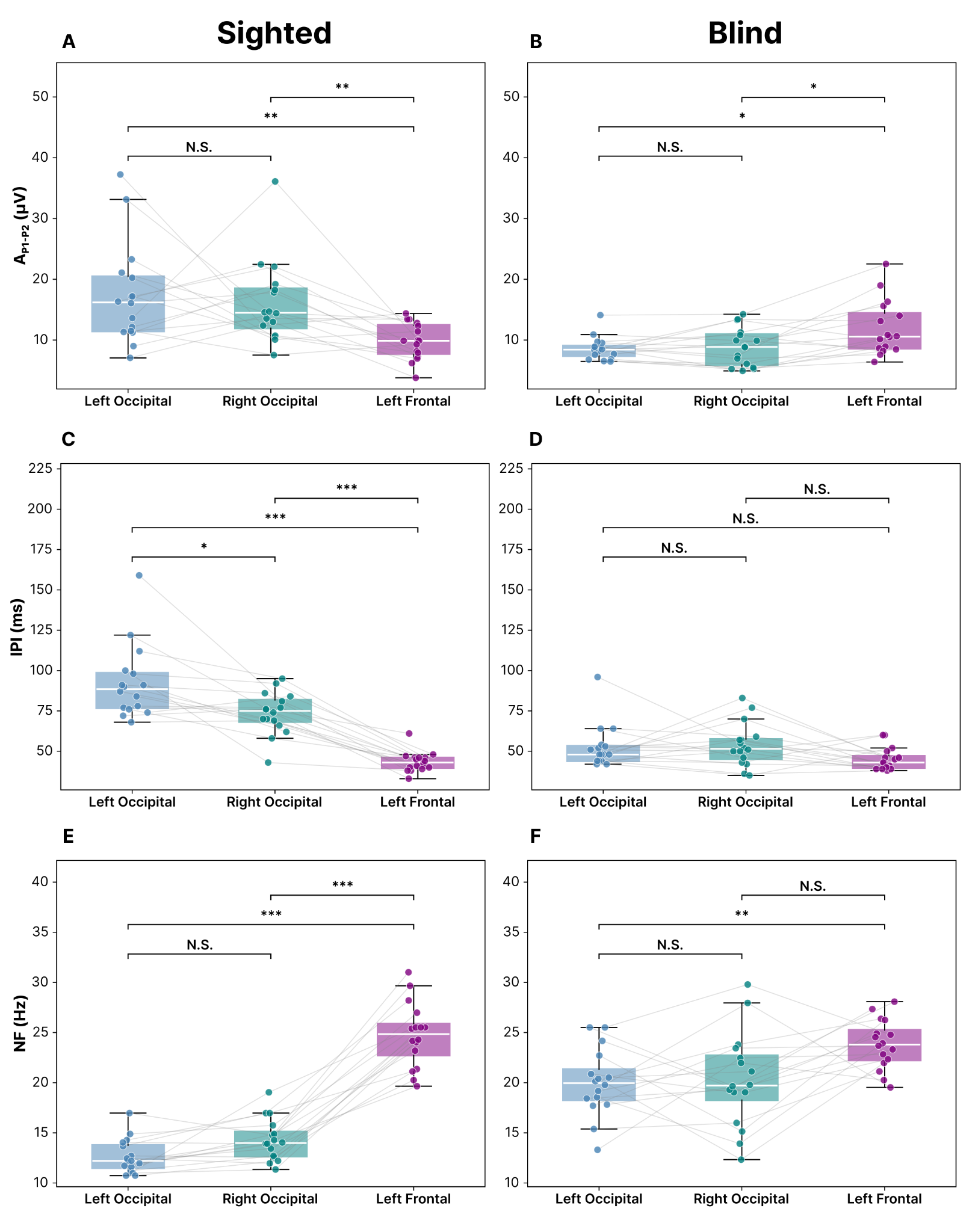

**Fig. S4. *Occipital and frontal oscillatory activity in sighted and blind individuals.*** Boxplots comparing left occipital (blue), right occipital (teal), and frontal (purple) cortex for A_P1-P2_, IPI and NF in sighted (A) and blind (B) individuals. Colored dots represent individual values; gray lines connect paired values across regions for the same individual. Differences across the three regions were assessed with a Friedman test for each metric and group, followed by pairwise Wilcoxon signed-rank tests, Bonferroni-corrected for multiple comparisons (ns = not significant; *p < 0.05; **p < 0.01; ***p < 0.001). In sighted individuals, the Friedman test was significant for all three metrics, and frontal stimulation elicited significantly higher NF values, lower IPI and A_P1-P2_ than both occipital sites, reflecting the canonical spectral differentiation between frontal and occipital circuits. This differentiation was attenuated in blind individuals, with several occipito-frontal comparisons failing to reach significance and with an opposite pattern concerning A_P1-P2_.

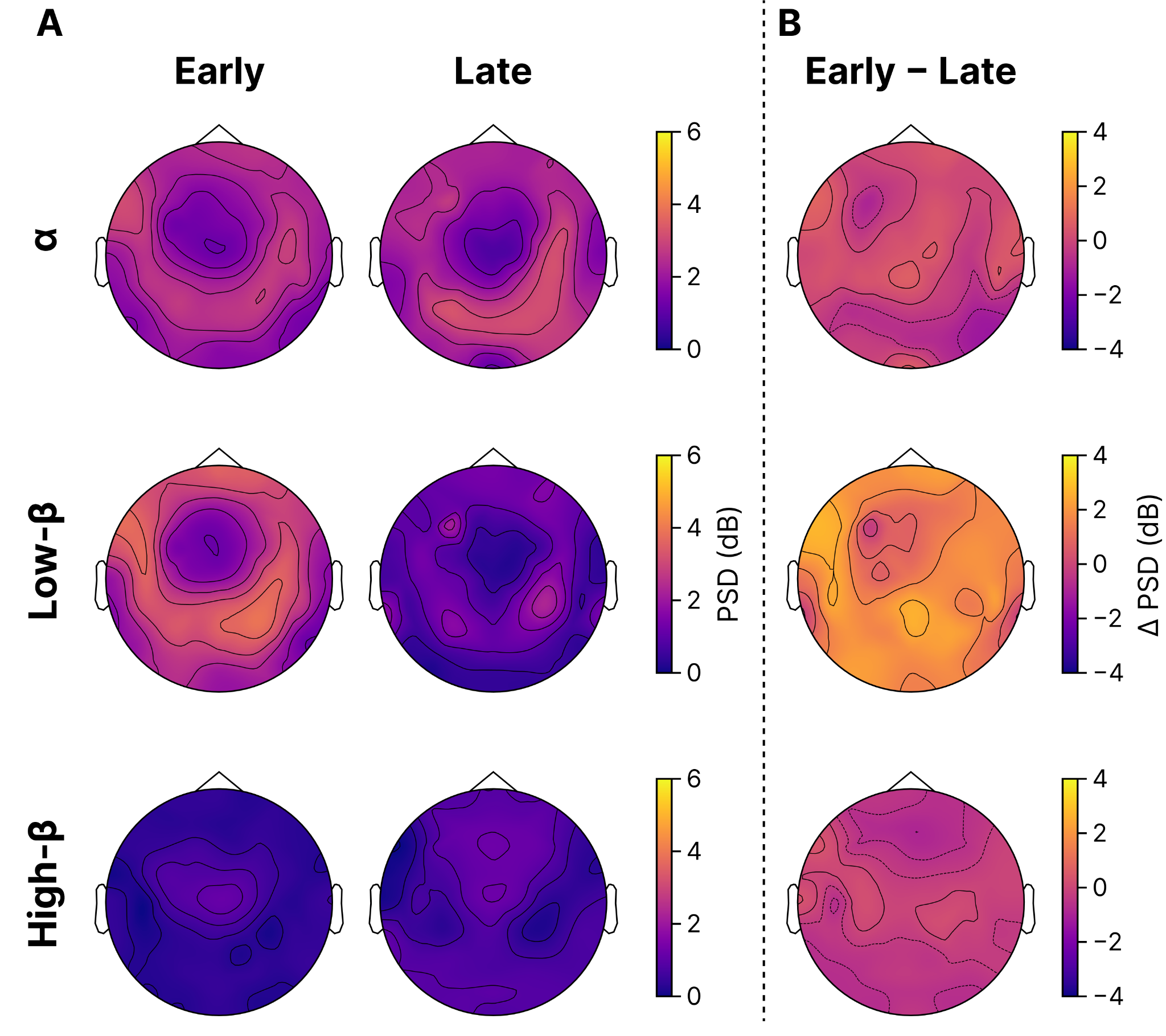

**Figure S5. *Topographical distribution in the alpha, low-beta, and high-beta bands for early onset blind individuals (Early) and late onset blind individuals (Late).*** Panel B shows the Early vs Late contrasts. No significant differences were found in any of the frequency bands.

**TMS-EEG preprocessing**

TMS-evoked potentials were preprocessed according to the following pipeline: (i) pulse artifact removal by replacing the signal from -2 to +5 ms with a mirrored and flipped version of the baseline between -2 and -9 ms, followed by a 4-ms-long moving average filter; (ii) detrending and high-pass filtering at 1 Hz with a zero-lag 3rd-order Butterworth filter; (iii) epoching between -600 and +600 ms around the TMS pulse; (iv) rejection of artifact-contaminated single trials and channels by visual inspection; (v) computation of the average reference and baseline correction by subtracting the mean pre-stimulus value; (vi) signal decomposition by ICA to remove components representing ocular and spontaneous muscle artifacts according to visual inspection; (vii) low-pass filtering at 45 Hz with a zero-lag 3rd order Butterworth filter and downsampling to 1000 Hz; (viii) interpolation of bad channels using spherical splines.

**TMS parameters**

We controlled for the spatial location and estimated intensity of the EF-max provided by the neuronavigation software in each stimulation site across blind and sighted individuals. For each TMS target, the distance between the single-subject EF-max coordinates of one group and the median EF-max coordinates of the other group was not significantly different (Mann-Whitney unpaired test) from the distance between the single-subject EF-max coordinates and the median EF-max coordinates of the same group. Thus, stimulation sites were comparable between the two groups, with median coordinates (in mm) of the EF-max in MNI reference space being [-16.6; 17.0; 58.3] and [-15.7; 14.1; 64.0] for the left frontal target, [-16.2; -85.1; 35.5] and [-14.7; -87.5; 29.7] for the left occipital target, and [17.4; -86.3; 25.3] and [16.5; -87.1; 32.3] for the right occipital target in blind and sighted individuals respectively.
